# New Mechanistic Insights into Prp5-mediated Prespliceosome Formation

**DOI:** 10.1101/2025.08.08.669338

**Authors:** Ching-Yang Kao, Wei-Yu Tsai, Yu-Lun Su, Che-Sheng Chung, Soo-Chen Cheng

**Author notes:** Center for Frontier Medicine, National Taiwan University Hospital, Taipei, Taiwan 100, Republic of China (Ching-Yang Kao). These authors contribute equally to this work (Ching-Yang Kao, Wei-Yu Tsai).

## Abstract

The spliceosome is a highly dynamic structure that undergoes continuous structural alterations through the sequential association and dissociation of small nuclear RNAs and protein factors during precursor mRNA splicing. These structural changes are driven by eight DExD/H-box RNA helicases that act at distinct stages of the splicing cycle. Among them, Prp5 and Sub2 are involved in prespliceosome formation, with Prp5 implicated in displacing the U2 snRNP component Cus2, and Sub2 in facilitating the release of the Msl5-Mud2 heterodimer. However, the precise mechanisms underlying the functions of these two proteins remain unclear. Here, we show that Sub2 is not essential for splicing *in vitro*, but it can enhance splicing independently of ATP. Strikingly, prespliceosome formation can proceed without ATP in the absence of either Sub2 or Cus2. Moreover, though ATP is required for prespliceosome formation under standard conditions, ATP hydrolysis is not. These findings reveal a coordinated interplay among Prp5, Sub2, Cus2 Mud2 and Msl5 during prespliceosome formation and indicate that ATP binding, rather than ATP hydrolysis, drives the early remodeling events that initiate spliceosome assembly.

## INTRODUCTION

Splicing of precursor mRNA (pre-mRNA) occurs within the spliceosome, a dynamic macromolecular complex that assembles through the sequential binding and release of its components on the pre-mRNA to align the 5’ splice site (5’SS) with the 3’ splice site (3’SS). The process begins with the binding of the U1 small ribonuclear protein particle (snRNP) to the 5’SS and of the dimeric protein complex Msl5-Mud2 to the branch site (BS), together forming the Commitment Complex (CC) (Séraphin and Rosbash 1989). Subsequently, Msl5-Mud2 is replaced by the U2 snRNP, forming the prespliceosome (PS), also known as complex A. Addition of the U4/U6.U5 tri-snRNP leads to the formation of a spliceosome complex containing all five snRNAs, referred to as the PreB complex, which subsequently undergoes structural rearrangements, releasing U1, forming the B complex, and U4 to form the active spliceosome, known as the B^act^ complex. The spliceosome can then catalyze two steps of transesterification, generating mature mRNA with the intron excised in a lariat structure. Upon completion of the splicing reaction, the mRNA is released from the spliceosome, which is subsequently dismantled to recycle its components.

The splicing process requires eight DExD/H-box RNA helicases, members of an RNA-dependent ATPase family. These helicases are generally thought to utilize the energy from ATP hydrolysis to drive structural changes in RNA molecules or ribonuclear protein complexes (Staley and Guthrie 1998; Cordin et al. 2012; Cordin and Beggs 2013; Liu and Cheng 2015; De Bortoli et al. 2020). Two DExD/H-box RNA helicases, Prp5 and Sub2, are involved in early steps of spliceosome assembly for formation of the prespliceosome. Prp5 is essential for recruitment of the U2 snRNP to the pre-mRNA (Perriman and Ares 2000). However, in the absence of the U2 component Cus2, ATP is dispensable for PS formation (Perriman and Ares 2000). Moreover, an ATP-binding mutant of Prp5 remains functional for cellular viability and *in vitro* splicing (Perriman and Ares 2000; Perriman et al. 2003; Liang and Cheng 2015), indicating that Prp5’s ATPase activity is primarily required to displace Cus2. It has been postulated that U2 adopts an inactive conformation in the presence of Cus2, with removal of Cus2 being necessary to activate U2 for its role in splicing (Perriman et al. 2003). Notably, Prp5 interacts with the spliceosome only transitorily, generally not being detected in association with it. However, it has been observed in spliceosomal complexes, specifically the Prp5-associated intermediate complex (FIC) and the PreA complex, which forms with BS-mutated pre-mRNAs that disrupt U2/BS base pairing (Liang and Cheng 2015; Plaschka et al. 2017), or with branchpoint A (brA)-deleted pre-mRNA (Zhang et al. 2021), respectively. Additionally, splicing in human extracts treated with the inhibitor spliceostatin A results in the accumulation of PreA-like complexes containing Prp5 (Zhang et al. 2024). These findings indicate that Prp5 must be released before the tri-snRNP can bind to the spliceosome, consistent with earlier biochemical evidence that Prp5 and the tri-snRNP are mutually exclusive on the spliceosome (Liang and Cheng 2015).

Similarly, although *SUB2* is essential for yeast viability, its deletion can be tolerated when combined with deletion of *MUD2* or certain *MSL5* mutations (Kistler and Guthrie 2001; Jacewicz et al. 2015). Genetic and biochemical evidence suggest that Sub2 plays a role in displacing Msl5-Mud2 to enable the binding of U2 to the BS during PS formation. When the interaction between Msl5 and the BS is weakened, either by Mud2 loss or Msl5 mutations, Sub2 becomes dispensable (Kistler and Guthrie 2001; Libri et al. 2001; Zhang and Green 2001). Nevertheless, the mechanisms underlying Sub2’s diverse functions remain unclear.

Sub2 is the yeast homolog of human UAP56, which was originally identified as a protein associated with U2AF (Fleckner et al. 1997), the human counterpart of Mud2. Apart from its role in splicing by facilitating PS formation, UAP56/Sub2 has been shown to play roles in transcription and nucleocytoplasmic export of mRNA (Gatfield et al. 2001; Jensen et al. 2001; Luo et al. 2001; Strässer and Hurt 2001; Strässer et al. 2002), as well as interacting with diverse protein partners to perform its various cellular functions (Zenklusen et al. 2002; Fidler and Ansari 2024). Nevertheless, the observation that Sub2 becomes dispensable when Mud2 is absent or when Msl5 function is impaired suggests that its primary role may lie in splicing.

To investigate the detailed mechanism underlying Sub2’s role in PS formation, we established a Sub2-depleted system for *in vitro* splicing analysis. Our results show that Sub2 is not essential for *in vitro* splicing but enhances splicing efficiency. Notably, in the absence of Sub2 or Cus2, PS formation does not require ATP, indicating that Cus2’s displacement can proceed independently of ATP. However, when both Sub2 and Cus2 are present, ATP is required, though not its hydrolysis. This observation aligns with previous findings in the human system, where ATPγS supports PS formation (Agafonov et al. 2016). Our data suggest that Sub2 may help stabilize Msl5 binding in conjunction with Mud2. Furthermore, the ATP requirement for PS formation appears to stem from Prp5, which requires ATP binding, but not hydrolysis, to induce a conformational change that enables it to displace Cus2, Msl5-Mud2, itself and possibly Sub2 as well.

## RESULT

### Sub2 is not essential for the in vitro splicing reaction

Previous studies have indicated that DExD/H-box RNA helicases Prp5 and Sub2 are involved in PS formation, which represents the first ATP-dependent step of the splicing process (Dalbadie-McFarland and Abelson 1990; O’Day et al. 1996; Fleckner et al. 1997). However, ATP becomes dispensable for PS formation in the absence of Cus2, and certain ATP-binding mutants of Prp5 retain splicing activity both *in vivo* and *in vitro* (Perriman and Ares 2000; Perriman et al. 2003; Liang and Cheng 2015). These findings indicate that Sub2’s role in PS formation may be ATP-independent.

To investigate the role of Sub2 in PS formation, we attempted to deplete Sub2 from splicing extracts for functional analysis. However, neither *in vitro* depletion using antibodies against Sub2 or its V5-tag nor *in vivo* depletion *via* the *GAL*-promoter system successfully eliminated splicing activity. Given that cells in which both *SUB2* and *MUD2* have been deleted remain viable, we pursued an alternative strategy by first generating a *MUD2*-deleted strain (ΔM), followed by subsequent deletion of *SUB2* to create a double mutant strain (ΔMS).

The ΔM extracts exhibited reduced splicing activity (Figure 1A, lane 2) and lower efficiency in PS formation, as revealed by immunoprecipitation of the splicing reactions performed in Prp8-depleted ΔM extracts using antibodies against the U2 component Lea1 (compare lane 9 with lane 5). However, we did not detect Msl5 binding to the pre-mRNA when splicing was performed under ATP-depleted conditions to arrest CC formation (Figure 1B, lane 10). Similarly, neither Lea1 nor Msl5 was observed on the BS mutant U257G pre-mRNA (Figure 1C, lanes 9 and 10), indicative of a failure in FIC formation. Since Msl5 and Mud2 form a stable dimeric complex, which only transiently associates with the pre-mRNA before PS formation, these results indicate that stable association of Msl5 with the pre-mRNA in the arrested CC and FIC complexes may depend on the presence of Mud2. Alternatively, the absence of Mud2 may affect Msl5 levels. Western blot analysis of splicing extracts confirmed that the Msl5 levels were greatly reduced in ΔM extracts (Figure 1D, lanes 3 and 4). However, Msl5 levels in total cell lysates were not affected by *MUD2* deletion (Figure S1), indicating that Msl5 becomes insoluble in the absence Mud2 and remains in the precipitate during splicing extract preparation. To determine if the low splicing activity in ΔM extracts was due to reduced levels of Msl5, we assessed if splicing activity could be restored with recombinant proteins. Expression of Msl5 alone in *Escherichia. coli* resulted in insoluble proteins. To overcome this issue, we co-expressed Msl5 and Mud2 in *E. coli*, and then purified the dimeric Mud2-Msl5 protein complex (M2M5) (Figure S2). Addition of M2M5 effectively restored the splicing activity of ΔM extracts (Figure 1E).

**Figure 1.**
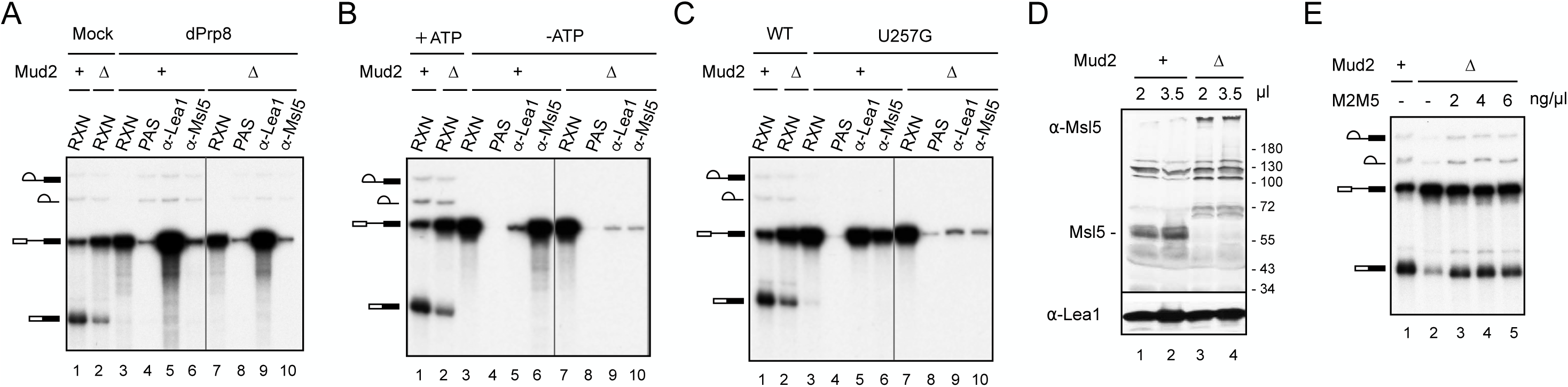
Characterization of ΔMud2 extracts. (A, B, C) Formation of the prespliceosome (A) the CC (B) and the FIC (C) in ΔMud2 extracts. Splicing was performed in wild-type (lanes 1 and 3-6) or ΔMud2 (lanes 2 and 7-10) extracts depleted of Prp8 (A) or ATP (B) using *ACT1* pre-mRNA, or in untreated extracts using U257G pre-mRNA (C), followed by precipitation with no (lanes 4 and 8), anti-Lea1 (lanes 5 and 9) or anti-Msl5 (lanes 6 and 10) antibody. RXN, 1/10 of the splicing reaction mixture; PAS, protein A-Sepharose; +, wild-type *MUD2*; Δ, *MUD2* deletion. (D) Western blot of wild-type (lanes 1 and 2) and ΔMud2 (lanes 3 and 4) extracts probed with anti-Msl5 and anti-Lea1 antibodies. +, wild-type *MUD2*; Δ, *MUD2* deletion. (E) Complementation of ΔMud2 extracts with purified recombinant M2M5. Splicing was performed in wild-type (lane 1) or ΔMud2 (lanes 2-5) extracts, to which various amounts of M2M5 was added. +, wild-type *MUD2*; Δ, *MUD2* deletion; M2M5, recombinant Mud2-Msl5 dimer.

Next, we deleted the *SUB2* gene to generate ΔMS cells. Spot assays demonstrated that whereas the ΔM cells exhibited slightly slower growth compared to wild-type cells, the ΔMS cells displayed a severe growth defect (Figure 2A). Western blot analysis revealed that, as in ΔM extracts, Msl5 levels were markedly reduced in the ΔMS extract (Figure 2B), leading to low splicing activity. However, adding recombinant M2M5 to the ΔMS extract substantially restored splicing activity (Figure 2C). These results indicate that although Sub2 is essential for cell viability, it is not required for splicing *in vitro*.

**Figure 2.**
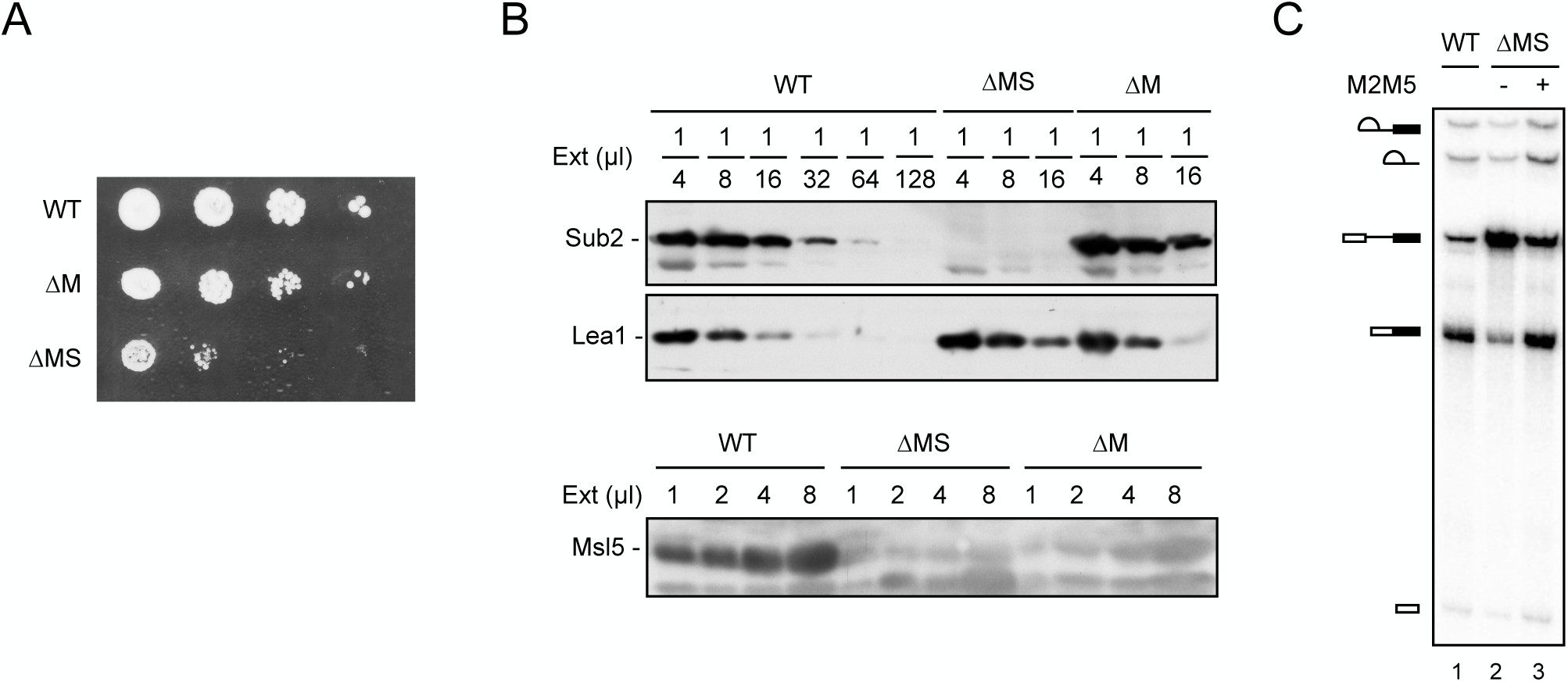
Characterization of ΔMS extracts. (A) Spot assays for growth at 30°C of serial dilutions of WT, ΔM and ΔMS cells. WT, wild-type; ΔM, *MUD2* deletion; ΔMS, *MUD2*, *SUB2*-double deletion. (B) Western blot of serial dilutions of WT, ΔM and ΔMS extracts, probed with anti-Sub2, anti-Lea1 and anti-Msl5 antibodies. WT, wild-type; ΔM, *MUD2* deletion; ΔMS, *MUD2*, *SUB2*-double deletion; Ext, splicing extract. (C) Splicing complementation of ΔMS extracts with purified recombinant M2M5 proteins. WT, wild-type; ΔMS, *MUD2*, *SUB2*-double deletion; M2M5, 2 ng/µl recombinant Mud2-Msl5 dimer.

### Complementary roles of Sub2 and Mud2 in promoting prespliceosome formation

Although Sub2 is not essential for splicing, it may enhance splicing efficiency—a role reminiscent of Prp22, which is dispensable for exon ligation but accelerates the reaction (Chung et al. 2025). To explore this potential function of Sub2, we conducted experiments using recombinant proteins. However, recombinant Sub2 proteins expressed and purified from *E. coli*. failed to improve splicing in ΔMS extracts, regardless of whether M2M5 was added or whether the HIS-tag was positioned at the N-or C-terminus of Sub2. Given that Sub2 is expressed in yeast at substantially higher levels than most splicing factors, over 50-fold higher than Prp2 or Prp22 (Figure S3), we instead used native Sub2 proteins by supplementing ΔMS extracts with diluted yeast extracts. To minimize the contribution of Msl5 and Mud2 from those extracts, we used ΔM extracts as the source of Sub2 and compared their complementation effects to those of ΔMS extracts, which lack Sub2 but contain similarly limiting levels of Msl5 as ΔM extracts.

Splicing reactions were carried out using ΔMS extracts complemented with increasing amounts of either ΔM (containing Sub2) or ΔMS extracts (lacking Sub2) at final protein concentrations ranging from 0.13 to 4 µg/µl (Figure 3A). Quantification of splicing efficiency revealed a modest enhancement upon addition of ΔM extracts, with less than a 10% increase at the highest amount added (4 µg/µl). In contrast, addition of diluted ΔMS extracts led to a decrease in splicing activity (Figure 3B). Although the reason for the inhibitory effect of the ΔMS extract remains unclear, it was consistently reproduced across three independent extract preparations. Nevertheless, our data indicate that Sub2 proteins present in the ΔM extract partially restored splicing activity in the ΔMS extract. This restorative ability was limited, likely due to the low levels of Msl5 in the ΔMS extract. Collectively, these results indicate that Sub2 can promote splicing under conditions where Msl5 is limiting.

**Figure 3.**
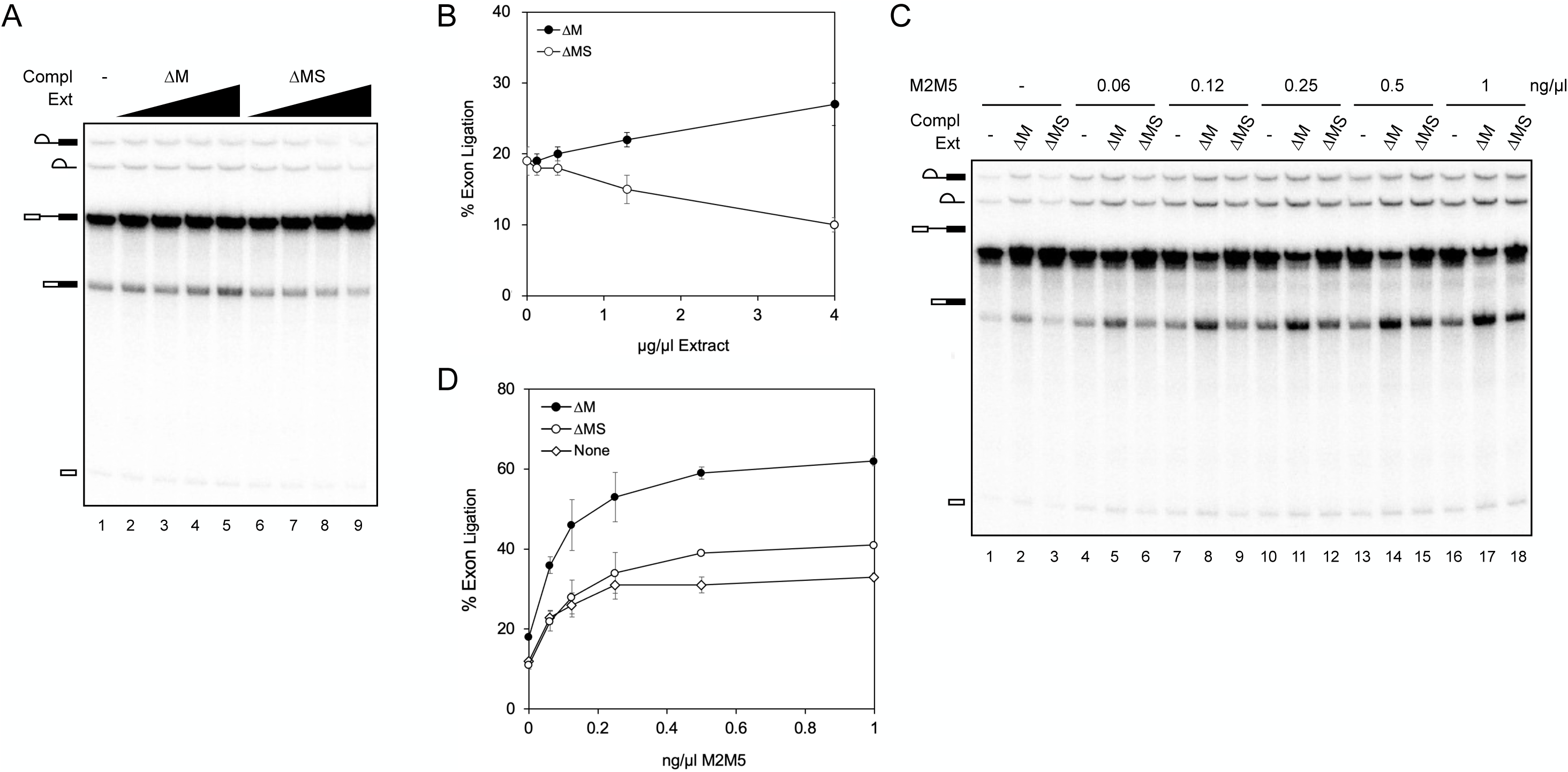
Splicing complementation of ΔMS extracts. (A) Splicing was performed in ΔMS extracts without (lane 1) or with the addition of various amounts of ΔM (lanes 2-5) or ΔMS (lanes 6-9) extracts. Lanes 2 and 6: 0.13 µg/µl; lanes 3 and 7: 0.4 µg/µl; lanes 4 and 8: 1.3 µg/µl; lanes 5 and 9: 4 µg/µl. Compl Ext, complementing extract; ΔM, *MUD2* deletion; ΔMS, *MUD2, SUB2*-double deletion. (B) The exon ligation efficiency of reactions from (A), calculated as the molar ratio of mRNA to the total of mRNA plus pre-mRNA. Data represent mean values from three independent experiments with standard deviation (SD) indicated. (C) Splicing was performed in ΔMS extract without (lanes 1, 4, 7, 10, 13 and 16) or with the addition of 2 µg/µl ΔM (lanes 2, 5, 8, 11, 14 and 17) or ΔMS (lanes 3, 6, 9, 12, 15 and 18) extracts, together with various amounts of M2M5 proteins. M2M5, recombinant Mud2-Msl5 dimer; Compl Ext, complementing extract; ΔM, *MUD2* deletion; ΔMS, *MUD2*, *SUB2*-double deletion. (D) The exon ligation efficiency of reactions from (C) was assessed from three independent experiments with SD indicated.

To assess if the complementation activity is additive, we supplemented ΔMS extracts with a low amount of ΔM or ΔMS extracts (2 µg/µl total proteins) in the presence of limiting concentrations of M2M5 (from 0.06 to 1 ng/µl) (Figure 3C). At all five M2M5 concentrations tested, the addition of ΔM extracts markedly enhanced splicing activity (lanes 5, 8, 11, 14 and 17), whereas ΔMS extracts had only a modest effect, primarily at higher M2M5 concentrations (lanes 6, 9, 12, 15 and 18). Quantification of splicing efficiency revealed a pronounced increase with increasing M2M5 levels up to 0.25 ng/µl. Reactions supplemented with ΔM consistently exhibited approximately 50% higher activity than those with ΔMS, which showed only slight improvement over reactions without added extracts (Figure 3D). These results suggest that Sub2 and Mud2 may function complementarily in PS formation, potentially by stabilizing Msl5 binding. This scenario supports previous findings that Sub2 exerts dual functions: promoting stable Msl5 binding to the pre-mRNA during CC formation, facilitating the release of Msl5 and Prp5 during the transition to PS formation (Kistler and Guthrie 2001; Libri et al. 2001; Zhang and Green 2001).

### Prp5’s function for PS formation is ATP-independent in the absence of Sub2

We reasoned that if Sub2 contributes to stabilizing Msl5 binding, then stable association of Msl5 with the CC complexes might be compromised in Sub2-depleted extracts. To test this possibility, we examined whether the CC2, the Msl5-containing CC, could be detected by immunoprecipitation of splicing reactions carried out in M2M5-supplemented ΔMS extracts (representing Sub2-depleted, or ΔSub2, extracts) under ATP-depleted conditions using anti-Msl5 antibody. Antibodies against Lea1, Prp5 and Prp8 were used as controls to detect U2, the FIC and U5, respectively. Precipitates were washed with standard buffers containing either 150 mM or 300 mM NaCl (Figure 4A). Our results show that while Lea1 and Prp8 remained associated with the pre-mRNA even after a high-salt wash, levels of Msl5-associated pre-mRNA were markedly reduced after washing with 300 mM NaCl buffer (compare lanes 3 and 9), indicating that an excess of recombinant Msl5-Mud2 complex may bind nonspecifically. Nonetheless, Msl5 was still detectable on the pre-mRNA in the absence of Sub2, indicating that CC2 formation does not require Sub2. Strikingly, both U2 (Lea1) and the tri-snRNP (Prp8), but not Prp5, associated with the pre-mRNA in a salt-resistant manner. Since U2 and Prp5 coexist in the FIC or the PreA complex, and prespliceosome formation involves release of Prp5 (Liang and Cheng 2015), the presence of Lea1 and absence of Prp5 indicate that the prespliceosome had formed. Given that addition of the tri-snRNP does not require ATP, our observation of Prp8 binding supports that the spliceosome had progressed to the PreB complex. Northern blot analysis of streptavidin-pulldown spliceosome confirmed the tri-snRNP association, further supporting this conclusion (Figure 4B). Together, these results indicate that splicing can progress to the PreB complex under ATP-depleted conditions even in the absence of Sub2, albeit less efficiently. This scenario indicates that Prp5 can promote PS formation and be released from the spliceosome without ATP when Sub2 is absent.

**Figure 4.**
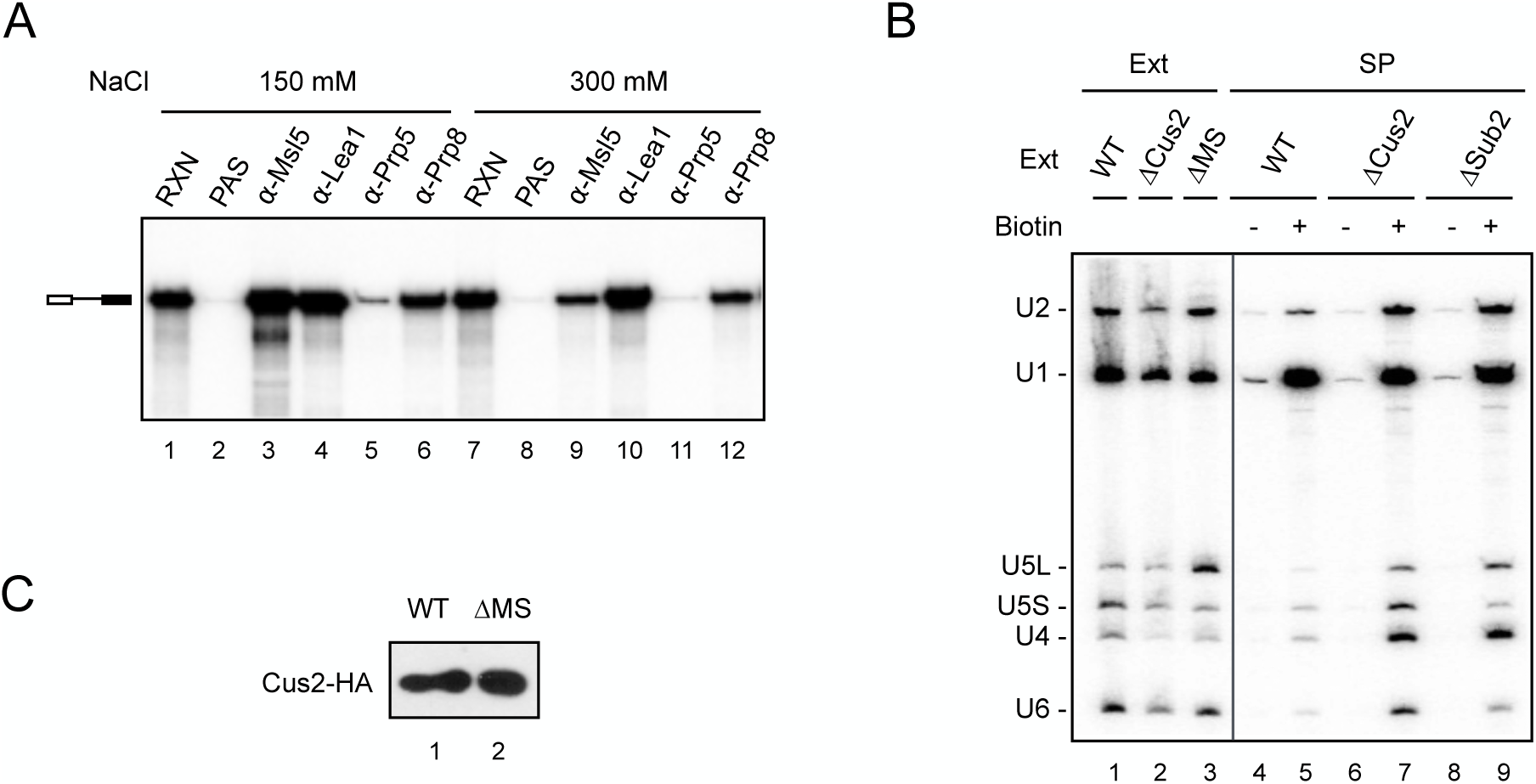
ATP is not essential for the formation of the Pre-B complex in Sub2-depleted extracts. (A) Splicing was performed in M2M5-supplemented ATP-depleted ΔMS extracts, and then the reaction mixtures were precipitated with no (lanes 2 and 8), anti-Msl5 (lanes 3 and 9), anti-Lea1 (lanes 4 and 10), anti-Prp5 (lanes 5 and 11) or anti-Prp8 (lanes 6 and 12) antibody. The precipitates were washed with standard buffer containing 150 mM or 300 mM NaCl. RXN, 1/10 of the splicing reaction mixture; PAS, protein A-Sepharose. (B) Northern blot of streptavidin-pulled down spliceosome. Splicing was performed pre-mRNA without (lanes 4, 6 and 8) or with (lanes 5, 7 and 9) biotin labels in WT (lanes 4 and 5), ΔCus2 (lanes 6 and 7) or ΔSub2 (lanes 8 and 9) extracts pre-incubated with glucose for 10 min. The reaction mixtures were precipitated with streptavidin agarose. RNAs from the precipitates and 2 µl of each extract (lanes 1-3) were analyzed by Northern blotting using probes for 5 snRNAs. Ext, splicing extract; SP, spliceosome; ΔMS, *MUD2*, *SUB2*-double deletion; M2M5, recombinant Mud2-Msl5 dimer. (C) Western blotting of Cus2-HA in wild-type and ΔMS extracts probed with anti-HA antibody following immunoprecipitation with anti-HA antibody. WT, wild-type; ΔMS, *MUD2*, *SUB2*-double deletion.

Prp5 has been implicated in displacing Cus2 in an ATP-dependent manner during PS formation (Perriman et al. 2003). Certain ATPase-deficient mutants remain functional in splicing when Cus2 is depleted (Liang and Cheng 2015). To rule out the possibility that Sub2 depletion affects Cus2 levels, we assessed Cus2 protein levels in ΔMS extracts. Cus2 was tagged with HA in both wild-type and ΔMS cells. Although Western blot analysis of splicing extracts showed weak protein signals, immunoprecipitation with anti-HA antibody to enrich Cus2 proteins clearly demonstrated comparable Cus2 levels in both the wild-type and the ΔMS extracts (Figure 4C). These results indicate that ATP is not required for PS formation when either Cus2 or Sub2 is depleted, and reveal that the ATP-dependent function of Prp5 extends beyond merely displacing Cus2.

### Prp5 mediates prespliceosome formation through ATP binding not ATP hydrolysis

Although DExD/H-box proteins are widely recognized for mediating structural changes in the spliceosome through ATP hydrolysis, studies on the human system have shown that ATPγS can support spliceosome assembly up to the stage preceding activation (Agafonov et al. 2016), indicating that ATP hydrolysis is not essential for PS formation. While this feature has not been reported in yeast, our findings support this scenario and indicate that Prp5’s function, previously associated with Cus2 displacement, may be independent of ATP hydrolysis. To test this possibility, we performed splicing in the presence of ATPγS and analyzed complex formation by immunoprecipitation with anti-Lea1, anti-Msl5 and anti-Prp8 antibodies (Figure 5A). Parallel reactions were conducted under normal and ATP-depleted conditions for comparison. Consistent with the observation of human spliceosomes, ATPγS permitted recruitment of U2 and the tri-snRNP (lanes 14 and 15), indicative of PreB complex formation. Under ATP-depleted conditions, Lea1 and Prp8 showed only a limited association with the pre-mRNA (lanes 9 and 10), whereas Msl5 accumulated more significantly (lane 8), likely reflecting retention in the CC. Notably, ATPγS reduced Msl5 level (lane 13) as the reaction progressed to formation of the PS and PreB complexes. These results indicate that, in yeast as in humans, formation of the PS and PreB complex does not require ATP hydrolysis.

**Figure 5.**
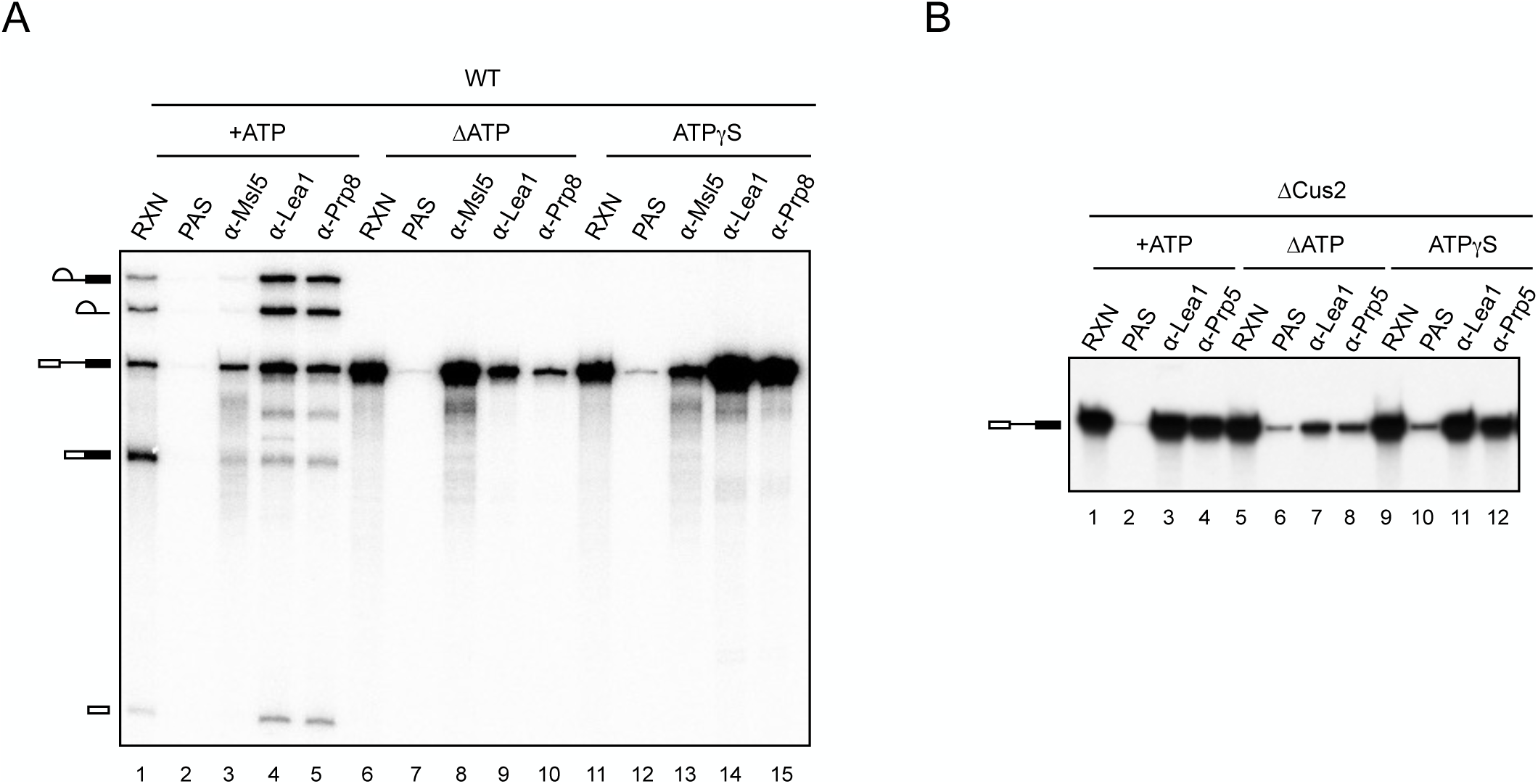
ATP hydrolysis is not essential for formation of the Pre-B complex or FIC. (A) Splicing was performed with wild-type *ACT1* pre-mRNA in wild-type extracts pre-incubated with 10 mM glucose (lanes 6-10), or in the presence of 2 mM ATP (lanes 1-5) or 2 mM ATPγS (lanes 11-15), followed by precipitation with anti-Msl5, anti-Lea1 or anti-Prp8 antibody. RXN, 1/10 of the splicing reaction mixture; PAS, protein A-Sepharose. (B) Splicing was performed with U257G mutant pre-mRNA in Δcus2 extracts pre-incubated with 10 mM glucose (lanes 5-8), or in the presence of 2 mM ATP (lanes 1-4) or 2 mM ATPγS (lanes 9-12), followed by precipitation with anti-Lea1 or anti-Prp5 antibody. RXN, 1/10 of splicing reaction mixture; PAS, protein A-Sepharose.

Our observations that displacement of Cus2 and Msl5 requires only ATP binding by Prp5 or Sub2 challenges the conventional view of energy requirement for disrupting protein-protein, protein-RNA, or RNA-RNA interactions. Moreover, it indicates that the ATP-bound forms of these DExD/H-box proteins represent their functional states. While Sub2 has not been shown to interact with pre-mRNA, Prp5 associates with pre-mRNA only transiently and is typically not detectable in assembled spliceosome complexes. However, Prp5 can be trapped at the FIC when U2/BS base pairing is impaired by BS mutations (Liang and Cheng 2015). To determine if ATP binding by wild-type Prp5 is required for FIC formation, we performed splicing with U257G mutant pre-mRNA in ΔCus2 extracts either depleted of ATP or supplemented with ATP or ATPγS followed by immunoprecipitation with anti-Lea1 and anti-Prp5 antibodies (Figure 5B). Our results reveal that both ATP and ATPγS supported binding of U2 and Prp5 to the pre-mRNA, whereas ATP depletion did not. Similar results were obtained for ΔS extracts, in which ATP or ATPγS stabilized the FIC (Figure S4). These findings suggest that although ATP is not required for PS formation in the absence of Cus2 or Sub2, stable association of wild-type Prp5 with pre-mRNA does require ATP binding. Thus, the ATP-bound form of Prp5 plays dual roles in PS formation – it stabilizes the intermediate complex, and promotes displacement of Cus2, Msl5 and Prp5 from the FIC.

## DISCUSSION

In this study, we investigated the functional role of Sub2 in splicing. Sub2 is a multifunctional protein involved in transcription and RNA transport in addition to its role in splicing. Its cellular abundance is at least 50-fold higher than that of most other splicing factors. Although *SUB2* is essential for cellular viability, the specific function underlying this essentiality remains unclear. A previous study used the *GAL*-promoter system to deplete Sub2 (Zhang and Green 2001), but we were unable to eliminate splicing activity through Sub2 depletion either *in vivo* or *in vitro*. Building on the previous observation that deleting *MUD2* can rescue the lethal growth phenotype of *SUB2* deletion, we generated a *mud2Δ*, *sub2Δ* double mutant yeast strain and prepared splicing extracts from these cells, which lacks Sub2 entirely. However, Msl5 was also severely depleted in the absence of Mud2, leading to low splicing activity in these extracts. Supplementation with recombinant Msl5-Mud2 dimer restored Msl5 function, yielding extracts selectively depleted of Sub2. Using these extracts, we found that splicing remains fully active in the absence of Sub2, indicating that Sub2 is not essential *per se* for the splicing reaction. This scenario contrasts with a previous report showing that CC2 formation is blocked in Sub2-depleted extracts (Zhang and Green 2001). One possible explanation for this discrepancy is that although Msl5 can transiently associate with pre-mRNA without Sub2, stable association may require Sub2, particularly when the spliceosome is stalled at the CC2 complex.

Although we show that Sub2 is not essential for the splicing reaction, we observed enhanced splicing efficiency in ΔMS extracts upon Sub2 addition. This suggests that Sub2 may help stabilize binding of Msl5 to the BS, potentially acting in a role similar to Mud2. This echoes previous genetic data indicating that Sub2 contributes to Msl5 stabilization (Kistler and Guthrie 2001). However, since we were unable to purify Msl5 alone for complementation, it remains unclear whether Sub2 plays a supportive role in splicing when Msl5 is abundant but Mud2 is absent.

The prior observation that *MUD2* deletion bypasses the requirement for *SUB2* led to the hypothesis that Sub2 may facilitate U2 recruitment by destabilizing Msl5-Mud2 binding to pre-mRNA, thereby enabling U2 positioning at the BS (Kistler and Guthrie 2001). Supporting this model, specific mutations in *MSL5* have also been shown to rescue the growth defect caused by *SUB2* deletion (Jacewicz et al. 2015). On the other hand, *MUD2* deletion or the individual suppressive *MSL5* mutations alone only elicit mild growth defects, which are exacerbated by *SUB*2 deletion. These findings indicate that when binding of Msl5 to the pre-mRNA is weakened, whether by Mud2 loss or Msl5 mutations, Sub2 plays a more critical role in stabilizing this transient interaction. Thus, our results support a model in which Sub2 plays dual roles in PS formation, by either stabilizing or destabilizing Msl5 binding depending on the cellular context. Such seemingly contradictory functions have been observed for other DExD/H-box proteins, such as Prp16 and Prp22, during the catalytic steps of splicing. Prp16 stabilizes Cwc25 binding to promote the branching reaction when mutations at the branchpoint impair Cwc25 association. Following branching, Prp16 facilitates Cwc25 removal from the catalytic center to allow 3’SS positioning (Chung et al. 2023). Likewise, Prp22 stabilizes Slu7 binding to facilitate the exon ligation reaction (Chung et al. 2025), and subsequently promotes mRNA release, together with Slu7 and itself, through ATP hydrolysis. Notably, the stabilizing roles of both Prp16 and Prp22 are ATP-independent, whereas their destabilizing activities require ATP hydrolysis. In contrast, neither the stabilizing nor the destabilizing function of Sub2 is dependent on ATP hydrolysis, indicating that Sub2 destabilizes Msl5 through a mechanism distinct from the canonical RNPase activity characteristic of other DExD/H-box proteins (Staley and Guthrie 1998; Schwer 2001).

Prp5 has been proposed to promote displacement of Cus2 from the U2 snRNP in an ATP-dependent manner during PS formation (Perriman and Ares 2000). Cus2 interacts with the U2 snRNP and modulates the structure of U2 snRNA (Yan et al. 1998; Rodgers et al. 2016). However, this interaction is relatively weak, particularly in yeast, making it difficult to detect the Cus2 association with U2 or with the spliceosome through immunoprecipitation or biochemical purification. It remains unclear whether Cus2 dissociates from the U2 snRNP before or after U2 binding to the pre-mRNA. In humans, the Cus2 homolog Tat-SF1 binds to U2 with higher affinity and it has been visualized in the cryo-EM structure of the human 17S U2 snRNP (Zhang et al. 2020). However, Tat-SF1 has not been detected in the PreA spliceosomal complex (Zhang et al. 2020; Tholen et al. 2022; Zhang et al. 2024), implying that it may be released or destabilized. In contrast, Prp5 has been detected in both the U2 snRNP and the PreA complex (Zhang et al. 2020; Tholen et al. 2022; Zhang et al. 2024), indicating that Prp5 remains associated with the spliceosome following Cus2 destabilization. Notably, ATP hydrolysis is not required for PS formation in either human or yeast extracts, implying that ATP binding by Prp5, rather than ATP hydrolysis, may be sufficient to induce a conformational change that facilitates Cus2 destabilization or release.

Strikingly, we found that in the absence of Sub2, ATP is dispensable for PS formation, although less than 10% of the pre-mRNA progressed to the PreB complex under these conditions. This suggests that Sub2 may help stabilize an intermediate complex, progression of which to the next stage requires Prp5 and ATP, consistent with complementation assays showing that Sub2 enhances splicing efficiency in the absence of Mud2. This phenotype resembles that observed in ΔCus2 extract and indicates that Cus2 may not be the sole target of Prp5’s activity during PS formation.

Previously, we demonstrated that mutations in the BS that disrupt U2-BS base pairing inhibit splicing by stalling the spliceosome prior to Prp5 release, resulting in formation of the FIC (Kao et al. 2021). Under these conditions, Prp5 is arrested on the spliceosome along with U2 and Msl5. UV crosslinking analysis revealed that neither U2 nor Msl5 binds to the mutated BS. Instead, Msl5 binds to an upstream cryptic BS within the *ACT1* intron, which contains a variation at the last nucleotide of the conserved BS sequence, while U2 dynamically interacts with sequences downstream of the BS. Restoration of proper U2-BS base pairing is essential for the reaction to proceed (Kao et al. 2021). A similar experiment performed using brA-deleted *ACT1* pre-mRNA also results in Prp5 arrest and formation of the PreA complex (Zhang et al. 2021). However, the cryo-EM structure of the PreA complex reveals formation of a U2-BS helix formed in the absence of the bulged branchpoint. The presence of Prp5 obstructs the necessary repositioning of the U1 and U2 snRNPs, thereby blocking subsequent tri-snRNP recruitment (Zhang et al. 2021). These findings suggest that formation of U2-BS helix alone is insufficient for Prp5 release. Instead, a specific RNA architecture with a bulged branchpoint may be crucial to trigger Prp5 release and enable PS formation. It remains uncertain if Sub2 or Cus2 associates with the FIC or the PreA complex, as their interactions with the pre-mRNA or U2 are weak and neither protein has been observed in structural studies. However, given that depletion of either protein bypasses the ATP requirement for PS formation and Prp5 release, it is likely that they are present on these complexes, potentially stabilizing binding of Msl5 to the pre-mRNA.

Msl5 recognizes and binds to the conserved BS sequence (Jacewicz et al. 2015), and this interaction can be detected in the splicing reaction through immunoprecipitation or UV crosslinking. However, Msl5 is not observed in the cryo-EM structure of the PreA complex (Zhang et al. 2021) or in the *in vitro* reconstituted early spliceosomal complex (E complex) (Li et al. 2019). Given that mutations at the branchpoint disrupt Msl5 binding (Jacewicz et al. 2015), it is possible that Msl5 binds to the upstream cryptic BS in the PreA complex assembled on brA-deleted *ACT1* pre-mRNA, similar to what has been observed in the FIC formed with BS mutated pre-mRNA (Kao et al. 2021). This interaction may be too weak or dynamic to be captured in cryo-EM structures. The FIC and PreA complex likely represent successive intermediates in prespliceosome formation, with the FIC preceding the PreA complex.

Based on current evidence, we propose a model (Figure 6) in which Msl5-Mud2, Sub2, Prp5 and Cus2 are all present in the FIC and/or PreA complex when U2 is recruited to the CC, binding to pre-mRNA sequences downstream of the BS. Msl5 associates stably with the BS in the presence of these components. Upon U2 engagement with the BS through base pairing, Msl5 is destabilized and released along with Mud2, Sub2 and Prp5, leading to formation of the prespliceosome. Cus2 may also be released at this stage. Since Sub2 is dispensable for splicing under standard conditions, Prp5 appears to be the sole essential DExD/H-box ATPase required for PS formation, potentially mediating displacement of Msl5-Mud2 and Prp5 itself. This displacement requires ATP binding, but not ATP hydrolysis. In the absence of Cus2, Mud2 or Sub2, Msl5 binding is weakened and can be disrupted even without ATP, although ATP binding by Prp5 can still promote the process. It remains unclear if Sub2 collaborates with Prp5 in this displacement activity. Our model does not fully explain how Mud2 depletion or Msl5 mutations can suppress the lethal phenotype associated with *SUB2* deletion, and the function of Sub2 that is essential to cell viability remains unknown.

**Figure 6.**
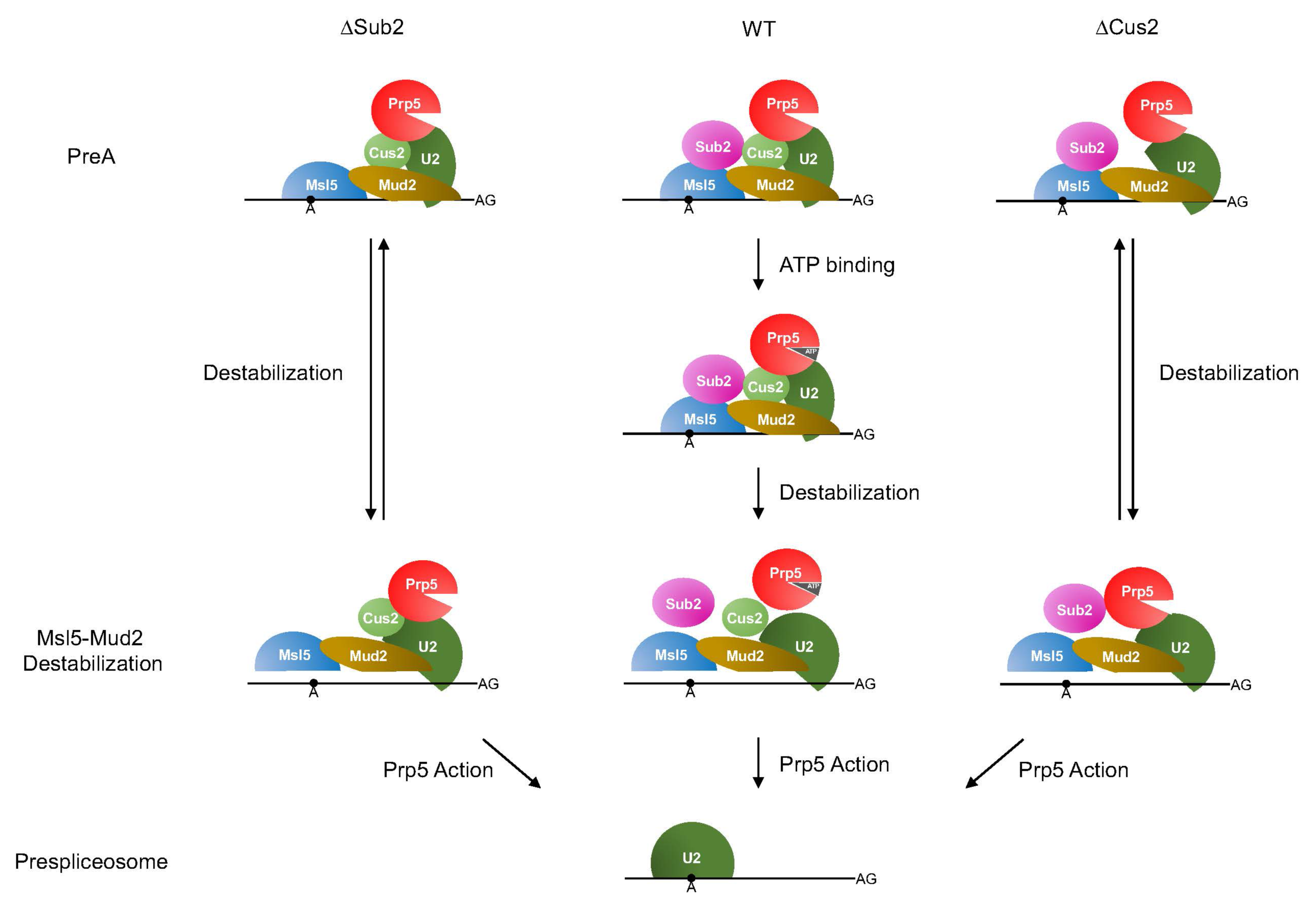
A model for the coordination of Sub2, Cus2, Mud2 and Prp5 in stabilizing and destabilizing binding of Msl5 on formation of the prespliceosome.

## MATERIALS AND METHODS

### Yeast strains

The yeast *Saccharomyces cerevisiae* strains used were BJ2168 (*MATa prc1 prb1 pep4 leu2 trp1 ura3*), YSCC028 (*MATa prc1 prb1 pep4 leu2 trp1 ura3 PRP5-V5 CUS2::LEU2*), YSCC029 (*MATa prc1 prb1 pep4 leu2 trp1 ura3 CUS2-HA*), YSCC043 (*MATa prc1 prb1 pep4 leu2 trp1 ura3 SUB2-4V5 MUD2::LEU2*), YSCC044 (*MATa prc1 prb1 pep4 leu2 trp1 ura3 SUB2::LEU2 pRS416.SUB2-4V5*), YSCC045 (*MATa prc1 prb1 pep4 leu2 trp1 ura3 MUD2::TRP1 SUB2::LEU2 pRS416.SUB2-4V5*), YSCC046 (*MATa prc1 prb1 pep4 leu2 trp1 ura3 MUD2::TRP1 SUB2::LEU2*) and YSCC047 (*MATa prc1 prb1 pep4 leu2 trp1 ura3 CUS2-HA MUD2::TRP1 SUB2::LEU2*)

### Antibodies and reagents

Anti-Lea1 antibody was generated by immunizing rabbits with full-length recombinant proteins. Anti-Msl5, anti-Prp5 and anti-Prp8 antibodies were raised by injecting rabbits with recombinant proteins comprising amino acid residues 273-412 for Msl5, 510-840 for Prp5, and 1-115 for Prp8. Protein A-Sepharose (PAS) was obtained from GE Healthcare Inc., Ni-NTA agarose was sourced from QIAGEN, and Proteinase K came from Cyrusbioscience Inc. SP6 RNA polymerase and RNasin were purchased from Promega.

### Construction of *SUB2*-4V5, *MUD2*-deletion and *MUD2*, *SUB2*-double deletion strains

To generate *SUB2-4V5* cells, a DNA fragment containing 534 base pairs (bp) of the *SUB2* 5’ upstream region, four tandem copies of the V5 epitope, and nucleotides 1-562 of the open reading frame (ORF) were cloned into the *BamH*I-*Xho*I site of pRS406 plasmid. The resulting plasmid was linearized with *Hpa*I and transformed into yeast strain BJ2168, followed by selection for *URA3* prototrophy. Selected transformants were grown in YPD medium and counter-selected for loss of *URA3* by plating on 5-fluoroorotic acid (5-FOA) plates, yielding *SUB2-4V5* cells. To create *MUD2*-deleted cells, the *MUD2* open reading frame (ORF) was replaced with a 2-kD DNA fragment of the *LEU2* cistron derived from YEP351 plasmid. To generate the *MUD2*, *SUB2*-double deletion cells, the *MUD2* ORF was first replaced with a 1.5-kilobase (kb) DNA fragment containing the *TRP1* cistron, amplified by polymerase chain reaction. The resulting Δ*MUD2* cells were then transformed with a pRS416-based plasmid carrying the *SUB2* cistron, which includes 0.85 kb of the promoter region, the ORF, and 0.1 kb of the 3’ untranslated region. Subsequently, the genomic copy of *SUB2* was deleted by replacing its ORF with a 2-kD *LEU2* cistron fragment, and the resulting cells were grown in YPD medium and counter-selected for loss of *URA3*-marked plasmid by plating on 5-FOA plates, yielding *MUD2*, *SUB2-*double deletion cells.

### Splicing extracts, substrates and reactions

Yeast whole-cell extracts and splicing reactions were prepared according to the method of Cheng *et al*. (Cheng et al. 1990). The pre-mRNA substrates were generated by *in vitro* transcription using SP6 RNA polymerase. *Eco*RI linearized pSP64-88 plasmid was used as the template to prepare standard actin substrate.

### Purification of recombinant Msl5-Mud2

A 1.6-kb DNA fragment encoding the HIS-tagged *MUD2* ORF and a 1.5-kb fragment of the *MSL5* ORF were cloned into plasmid pRSDuet-1 at the *Nco*I-*Bam*HI and *Aat*II-*Xho*I sites, respectively. The resulting plasmid was transformed into *E. coli* Rosetta cells for protein expression. Transformants were cultured in 1 liter of LB broth containing 50 µg/ml kanamycin until reaching an optical density (OD_600_) of 0.5. Protein expression was induced with 1 mM isopropyl β-D-1-thiogalactopyranoside and continued at 18°C for 4 hours. Cells were harvested, resuspended in 20 ml PBS, and lysed using a Microfludizer at 8,000-10,000 psi for 2-4 cycles. Following centrifugation to remove cell debris, the lysate was transferred to 50 mM Na_2_HPO_4_ (pH 8.0), 0.3 M NaCl and 10 mM Imidazole, and Msl5-Mud2 dimer was purified using a 5-ml Ni-NTA agarose column.

### Immunoprecipitation and immunodepletion

Immunoprecipitation of spliceosomes was performed as described previously (Tarn et al. 1993). For each 20 µl of splicing reaction mixture, 10 µl of protein A-Sepharose (PAS) conjugated with 5 µl of anti-Lea1, 5 µl of anti-Mls5, 12 µl of anti-Prp5 or 1 µl of anti-Prp8 serum was used. To deplete Prp5 and Prp8, each 100 µl of the cell extract was incubated with 100 µl of the anti-Prp5 or anti-Prp8 antiserum conjugated to 50 µl of PAS.

## SUPPLEMENTAL MATERIALS

Supplemental material is available for this article.

## ACKNOWLEDGMENTS

We thank J.M.C. Danac for generating the Msl5-Mud2 co-expression system, T.-H. Wu and H. L. Wai for technical assistance, and J. O’Brien for English editing. This work was supported by grants from Academia Sinica and National Science and Technology Council (Taiwan) (111-2311-B-001-008-MY3).

